## Supplemental figures and tables for "Structure of MlaFB uncovers novel mechanisms of ABC transporter regulation"

### Supplementary figures

**Supplemental Figure 1. Uncropped complementation plates and gels corresponding to Figures 1B, 1D, 4C, 5K and 5L.**

**A** **Phenotype of MlaF and MlaB, full image from Fig 1B**

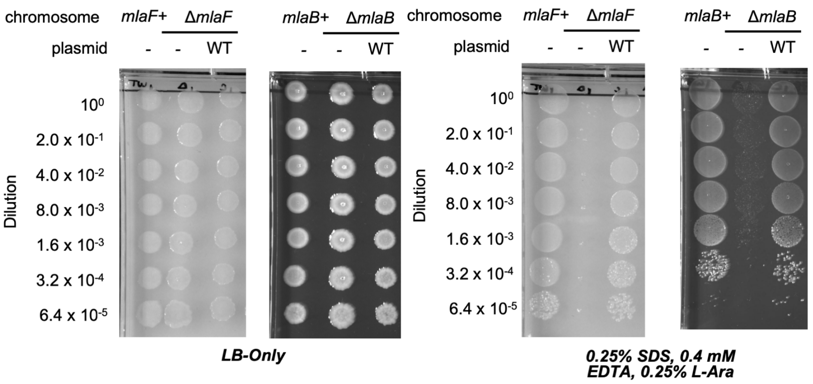


**B** **Gel of purified MlaFB Full gel from Fig 1D**

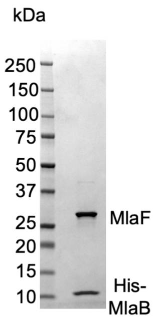


**C** **Pull downs: Walker mutants, CTD mutants and deletion, full image from Fig 4C**

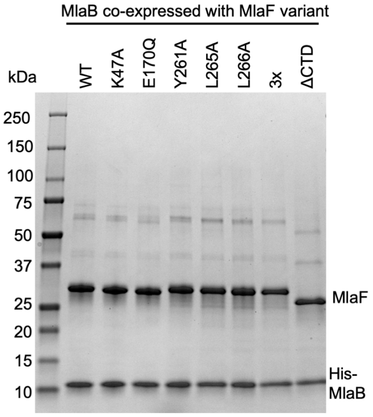


**D** **Full gel from Fig 5K**

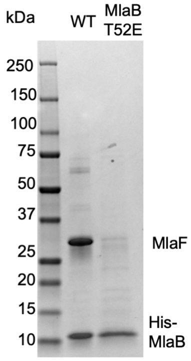


**E** **Full gel from Fig 5L**

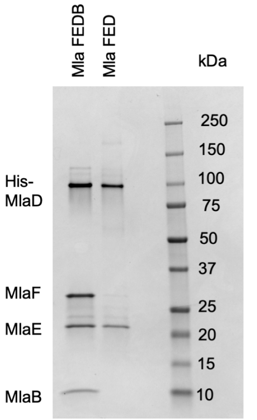

**Supplemental Figure 2. Optimization of conditions for growth analysis on different agar/media.** The concentrations of SDS and EDTA for different agar/media that resulted in optimal conditions for the assay are highlighted in blue.

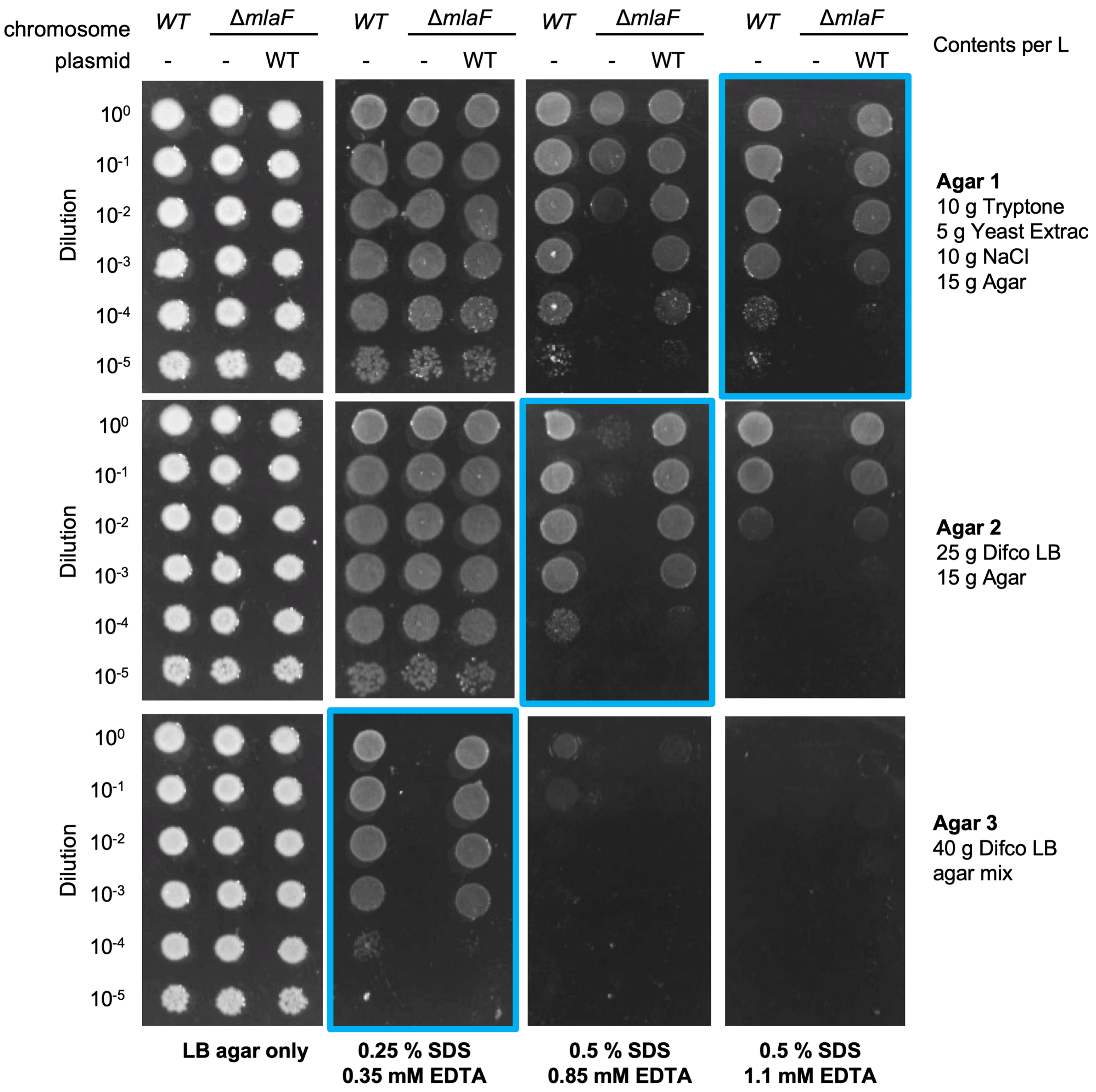

**Supplemental Figure 3. Position of Q-loop and subdomain conformations in different structures of MlaF.** *(A)* Docking of ADP-bound $MlaF_2B_2$ complex into previous low-resolution EM density map of *E. coli* MlaFEDB (EMD-8610). *(B)* Comparison between our two crystal structures (ADP-bound and apo), aligned on the catalytic subdomain. *(C)* Q-loop residues Gln92 (purple) from the ADP-bound MlaF structure and Gln89 from the ATP$\gamma$S-bound MetNI structure (pink; PDB: 6CVL[86]), shown as sticks.

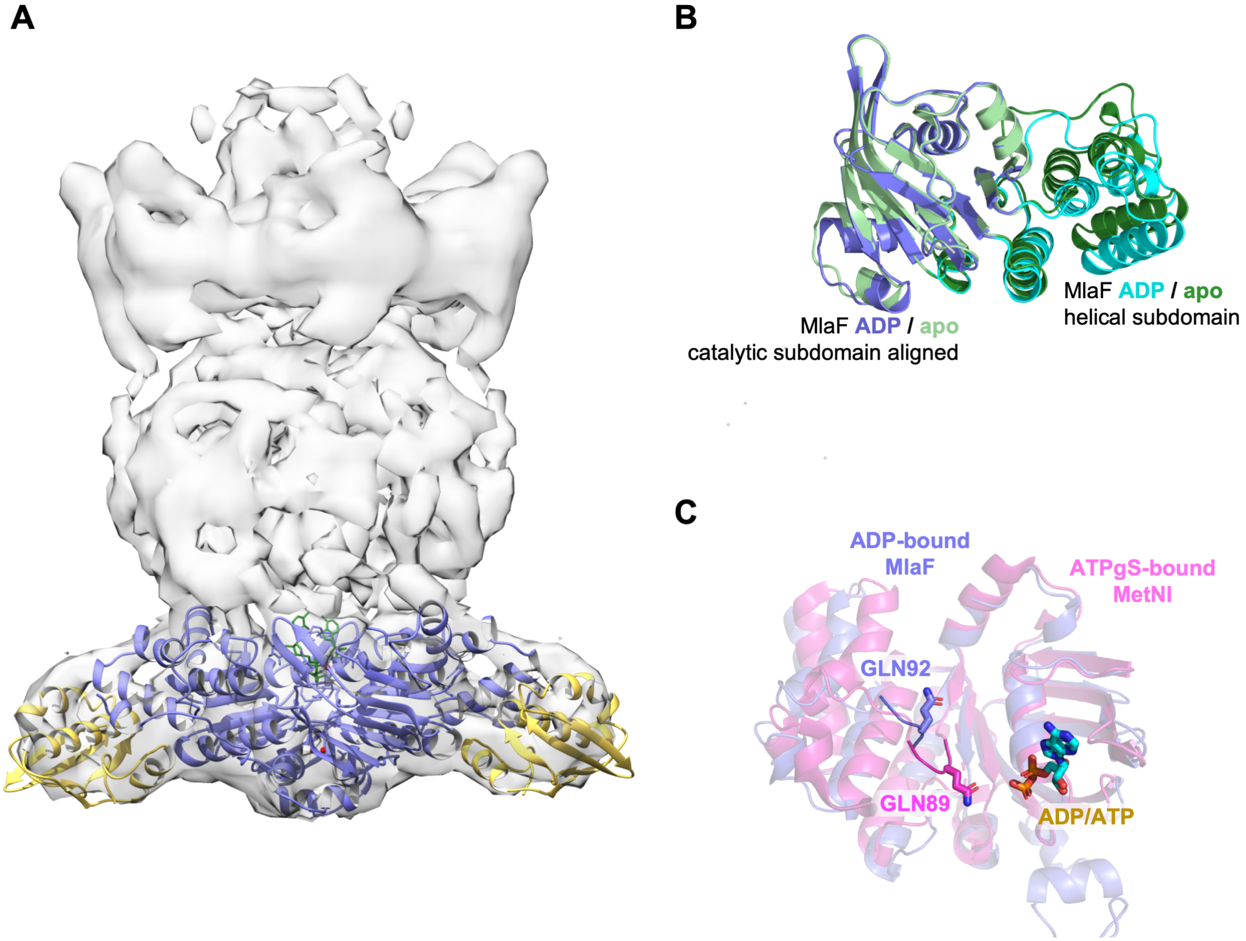

**Supplemental Figure 4. Sequence alignment of Mla, TGD-like MCE systems, o-antigen and teichoic acid transporter.**
The conserved LSGGQ/M motif is highlighted in green

```
MlaF E. coli                        ----------------MEQSVANLVDMRDVSFTRGNRCIFDNISLTVPRGKITAIMGPSG   44
MlaF R. rickettsii                  -----------------MSVEEFKIKIRSLYKSFANHKVLDGIDLDVKKGSSLVILGGSG   43
MlaF W. pipientis                   -------------------MCNPVISISDLCLSFYNRVVLNHLNLNISRGESLAILGGSG   41
MlaF E. ruminantium                 ---------------------MYCISLSDLHIAFHDHKILNGINLNVAYGESLVILGESG   39
MlaF N. moscoviensis                -----------------------MLKLVGVKKRLGGQPVLQGIDLTIPPGKLTTIIGRSG   37
MlaF F. tularensis                  ---------------------MCDISFKNVSFYRDSRCIYDDITFTIPTNKITTILGPSG   39
MlaF N. gonorrhoeae                 ------------------MSPSPFIEMKDVAFAYGDRPILNDINFSIPQGNFAAVMGGSG   42
MlaF B. cepacia                     ---------------MSPTPTETLLELRDVDFGYGDRLVLSNLNLRFGRGQVVAVMGGSG   45
MlaF P. aeruginosa                  ----------------MSTDSAYAVELKGLTFKRGSRAIFDNIDVRIPRGKVTGIMGPSG   44
MlaF L. pneumophila                 -------------------MPDNWVEIKNLFFSLDDRCIFDGIDMVVKQGKITAIMGPSG   41
MlaF A. baumannii                   ------------MNNKTPLSTQSLIEVKNLSFNRGERVIYDNISLNIRRGQITAIMGPSG   48
MlaF P. luteoviolacea               ---------------MESDLSQSIVEVKDVTFSRGERVIYKNMSFSVPRGKITAIMGPSG   45
MlaF H. influenzae                  -------------------MNQNLIEVKNLTFKRGDRVIYDNLNLQVKKGKITAIMGPSG   41
MlaF V. cholerae                    ------------------MSQSDLVTIKNLRFSRSQRVIFDDIDLHVPKGKVTAIMGPSG   42
MlaF K. pneumoniae                  ----MLARQVFSKDDFMSQTQANLVEVRGIRFSRGDRVIFDDISLSVPRGKITAIMGPSG   56
Teichoic Acid Exporter A. herbarius MEHAVIVENVTKKYKLFKRTSERLLDMILPGGYGEDFYALRNVSFTADKGDVIGIVGVNG   60
O-Antigen Exporter A. aeolicus      --MMIRVFDVWKKYKYYKKPQDRLKEIIFRKPFHEELWVLKGINLEIEKGEVLGIVGPNG   58
Maltose Importer E. coli            ---------------------MASVQLQNVTKAWGEVVVSKDINLDIHEGEFVVFVGPSG   39
                                                              .               : .    ..   .:* .*

MlaF E. coli                        IGKTTLLRLIGGQIAPDHGEILFDGENIPAMSR-SRLYTVRKRMSMLFQSGALFTDMNVF   103
MlaF R. rickettsii                  SGKSVLIKNIVGLIKPDKGKIFIDNVEIQDISS-KQKFEMMDGIGFLFQGGALFDSLNIR   102
MlaF W. pipientis                   SGKSVLAKTIIGLLLPDSGSIHV--------NS------KSKSFGVLFQNSALFDYITVW   87
MlaF E. ruminantium                 SGKSVLTKTILGLISPVSGKVIVDGIDVKHNKS------LSKNFGVLFQNCALFDSLTVC   93
MlaF N. moscoviensis                EGKSVLLKHMIGLLQPDEGQVWVDGVEISRLRG-HALNEVRKRFAMLFQGAALFDSMTVF   96
MlaF F. tularensis                  AGKTTILQLIAGLIKPTLGKICIDNAIIEKNTKETQLEKLRHRMGFLFQSGALFTHLSVY   99
MlaF N. gonorrhoeae                 SGKTTLMRLITGQIRPQSGQVLIEGRDLAGFSA-DELYEHRRRMGVLFQHGALFTDLSVF   101
MlaF B. cepacia                     CGKTTVLRLIGGLVRARRGQVLFDGADVGAQTR-DGLYALRRKMGMLFQFGALFTDMSVF   104
MlaF P. aeruginosa                  CGKTTLLRLIASQLRPSKGEVWVNGQNLPQLSR-GDLFDMRKQFGVLFQSGALFTDLDVF   103
MlaF L. pneumophila                 SGKTTLLRLIGAQLTPDSGQIYVNGQNIHQLSR-KALYETRRNMGLLFQSSALFTHLSVF   100
MlaF A. baumannii                   TGKTTLLRLIGGQLVPDHGEVLLDGKDIAQMSR-QELFAARARMGMLFQSGALFTDMSVY   107
MlaF P. luteoviolacea               IGKTTMLRLIGGQLRPDSGDITFMGSSVPAMSR-SELYQARKQMSMLFQSGALFTDMSVF   104
MlaF H. influenzae                  IGKTTLLKLIGGQLMPEQGEILFDGQDICRLSN-RELYEVRKRMGMLFQSGALFTDISTF   100
MlaF V. cholerae                    IGKTTLLRLIGGQLLPEQGEIWFDGENIPTLSR-RKLYRARKKMSMLFQSGALFTDLNVF   101
MlaF K. pneumoniae                  IGKTTLLRLIGGQIPPDRGEILFDGENVPQMTR-SRLYTVRKRMSMLFQSGALFTDLNVF   115
Teichoic Acid Exporter A. herbarius SGKSTLSNIIAGILPPTSGTIKIDGQ-ASLIAI----------------SSGLNNQLTGR   103
O-Antigen Exporter A. aeolicus      AGKSTLLKVITGVTEPDKGFVERSGKVVGLLEL----------------GTGFNYELSGL   102
Maltose Importer E. coli            CGKSTLLRMIAGLETITSGDLFIGEKRMNDTPP------AERGVGMVFQSYALYPHLSVA   93
                                     **:.: . : .      * :                              .:   :

MlaF E. coli                        DNVAYPLREHTQLPAPLLHSTVMMKLEAVGLRG-AAKLMPSELSGGMARRAALARAIALE   162
MlaF R. rickettsii                  DNITFETK---NLSKKEKNDLAGAKLNAVGLSPRILELYPAELSGGMQKRVALARAICST   159
MlaF W. pipientis                   ENISFNYKKRFNIDKKKAKQLAIEKLNSVGLEENISDMFPIELSGGMKKRVALARAIAHN   147
MlaF E. ruminantium                 ENITFNFHYRFNCNKNSAREIAASGLNLVGLSNDVLDRYPIELSGGMKKRIALARAIVSE   153
MlaF N. moscoviensis                ENVAFPLRERLRMKGAEVTKRVDEKLAQVGLGG-MGHKFPAELSGGMRKRAGLARALVMQ   155
MlaF F. tularensis                  DNIAFPLRKNTNLDEKLIRNIVLLKLQAVGLAH-TINMMPSELSGGMARRVALARSIAMD   158
MlaF N. gonorrhoeae                 DNIAFPMRELTQLPEAVIRDLVLLKLNAVGLRG-VENLMPSELSGGMSRRVALARTIALD   160
MlaF B. cepacia                     ENVAFALREHTDLPEDLIRDLVLMKLNAVGLRG-ARDLMPSEVSGGMARRIALARAIALD   163
MlaF P. aeruginosa                  ENVAFPLRVHTQLPEEMIRDIVLMKLQAVGLRG-AVELMPDELSGGMKRRVALARAIALD   162
MlaF L. pneumophila                 ENVAFPLREHTELNASMIRNIVLMKLEAVGLRG-AAHLMPAQLSGGMCRRVALARTIALD   159
MlaF A. baumannii                   ENVAFPIRAHTKLSENLIAELVALKLESVGLRG-TEQLMPTELSGGMNRRVALARAIALD   166
MlaF P. luteoviolacea               DNVAFPLREHTQLSEDLIRLVVLMKLQAVGLRG-ARELMPAELSGGMARRAALARAIALD   163
MlaF H. influenzae                  DNVAFPIREHTHLPENLIRQIVLMKLEAVGLRG-AAALMPSELSGGMARRAALARAIALD   159
MlaF V. cholerae                    DNVAYPLREHTELDEAMIKTLVLLKLEAVGLRG-AAYLMPSELSGGMARRAALARAIALD   160
MlaF K. pneumoniae                  DNVAYPLREHTHLPPALLHTTVMMKLEAVGLRG-AAKLMPSELSGGMARRAALARAIALE   174
Teichoic Acid Exporter A. herbarius ENIELKCLM-LGFSKKQIRAMEPDIIEFADIGK-FIDQPVKTYSSGMKSRLGFAISVNID   161
O-Antigen Exporter A. aeolicus      ENIYVNASL-LGLSRREIDEKLESIIEFSELDD-FINKPLKTYSSGMIMRLAFSIAIHTE   160
Maltose Importer E. coli            ENMSFGLKLAGA-KKEVINQRVNQVAEVLQLAH-LLDRKPKALSGGQRQRVAIGRTLVAE   151
                                    :*:                           :            *.*   * .:. ::

MlaF E. coli                        PDLIMFDEPFVGQDPITMGVLVKLISELNSALGVTCVVVSHDVPEVLSIADHAWILADKK   222
MlaF R. rickettsii                  PSILFLDEPTTGLDPIMANVINELIIKIQEELEATTITITHDMISAAKIAKEVAMIYQGK   219
MlaF W. pipientis                   PEIIILDEPTSGLDPIMSNIVNEIIMSLSKNLHSTIVTITHDIYSTFRIANKIAVLYEGN   207
MlaF E. ruminantium                 PKIIILDEPTSGLDPIMSDIVNDIIVKCHKKFKLTIITITHDISSAFKIASKIAVLYAGK   213
MlaF N. moscoviensis                PEIILFDEPTTGLDPLMAKSIHDLITSTQRTFGFTAVMVSHEIPEIFGISDYVAMLKQGK   215
MlaF F. tularensis                  PDIMMYDEPFTGQDPASFNKLLELISTLNESLNMTSIIVSHDIQESLSISDHIIIVGNKK   218
MlaF N. gonorrhoeae                 PEIMLYDEPFTGLDPISLGVIAHLISRVNKALRSTSIMVTHDIEKSLEIVDQVIFLAHGE   220
MlaF B. cepacia                     PQLIMYDEPFAGLDPISLGITANLIRTLNEALGATSILVTHDVPESFAIADYVYFLANGG   223
MlaF P. aeruginosa                  PQILLYDEPFVGQDPIAMGVLVRLIRLLNDALGITSIVVSHDLAETASIADYIYIVGDGR   222
MlaF L. pneumophila                 PVLMMYDEPFTGQDPISMGVLVRLIQRLNQLLHTTTIIVSHDVEETCSIADYIYLISGGK   219
MlaF A. baumannii                   PDLIMYDEPFAGQDPIVKGVLTRLIRSLREALDLTTIIVSHDVPETLSIADYIYVVAEGK   226
MlaF P. luteoviolacea               PELIMYDEPFAGQDPISMGVLVRMIKSLNEVLRLSSLIVTHDVTEVMSIADHVVIIAEQG   223
MlaF H. influenzae                  PDLIMFDEPFTGQDPISMGVILSLIKRLNEALNLTSIVVSHDVEEVLSIADYAYIIADQK   219
MlaF V. cholerae                    PELIMYDEPFVGQDPITMGVLVELIRNLNRALGVTSVVVSHDVPEVMSIADWVYLLADGK   220
MlaF K. pneumoniae                  PDLIMFDEPFVGQDPITMGVLVKLISELNSTLGVTCIVVSHDVPEVLSIADYAYIVADKK   234
Teichoic Acid Exporter A. herbarius PDVLVIDEALSVGDQTFADKCLDKMN-EFKERGKTIFFISHSIGQVKSFCEKALWLEYGE   220
O-Antigen Exporter A. aeolicus      PECFIIDQALAVGDAHFQQKCFRKLK-EHKQKGGSIIFVSHDMNAVKILCDRAILLHKGE   219
Maltose Importer E. coli            PSVFLLDEPLSNLDAALRVQMRIEISRLHKRLGRTMIYVTHDQVEAMTLADKIVVLDAGR   211
                                    *  :. *:     *          :         : . ::*.      : .    :

MlaF E. coli                        IVAHGSAQALQANPDPRVRQFLDGIADGPVPFRYPAGDYH-ADLLPGS-----          269
MlaF R. rickettsii                  IKWYGSKDDMRNSDNPYLKQFINGLTTGPIEV---------------------          251
MlaF W. pipientis                   IIFHGTIKEIQNSNNEYIKKFIAPVQFMS------------------------          236
MlaF E. ruminantium                 IIVCESVQEIKNTDNQYVRNFIHRNY---------------------------          239
MlaF N. moscoviensis                IAAMAPPAEFQRTADPEITEFISVGGTMALPAAPPNR----------------          252
MlaF F. tularensis                  IIASDSPENIKNSKDQQIQNFLAGK---PLDYNYHNHNDL-EKNFFKKEILGS          267
MlaF N. gonorrhoeae                 IMFSGSPQEMRELDSPWVRQFVGGLADGPVAYRYPAQTSL-QQDLLG------          266
MlaF B. cepacia                     VLAQGTPDELRASTDPSVRQFIDGAPDGPFKFHYTSPPLA-ADFGLGGGR-A-          273
MlaF P. aeruginosa                  VLGHGTPDVLKETDDPRIRQFVKGIPDGPVPFHYPARDYR-ADLLGER-----          269
MlaF L. pneumophila                 IIGQGTPEELKRSIEPQIRQFMHGEADGVVPFHYPAKPYT-EELLNA------          265
MlaF A. baumannii                   IQGEGTPEELQAYASPFVKQFLTGSAEGPVEYQFSHQAYL-DNEVRP------          272
MlaF P. luteoviolacea               VIGSGSPEQMLAHSSPLVQQFLQGLSDGPVPFHFKAAPYE-EELLDGGV----          271
MlaF H. influenzae                  VIAEGTSEQLLQSQDLRVVQFLKGESDGPVRFKYPAQDYV-KELFE-------          264
MlaF V. cholerae                    VIAQGSPQALRDNPDPRVQQFLCGDADGPVPFRFPAQPIE-QELFSAK-----          267
MlaF K. pneumoniae                  IVAHGSAQSLRENTDPRVRQFIDGIADGPVPFRYPAGDYH-HDLLGIGS----          282
Teichoic Acid Exporter A. herbarius VRGYGTVAEIIPQYEKFLKEYRAMSDKEKRQYKERVMRKQQGEFLQAAVK---          270
O-Antigen Exporter A. aeolicus      IIEEGSPETVTQAYYKL------------------------------------          236
Maltose Importer E. coli            VAQVGKPLELYHYPAD-------------------------------------          227
                                    .
```

**Supplemental Figure 5. Comparison of our *E. coli* MlaFB crystal structure (2.60 Å resolution) and coordinates deposited for *A. baumannii* MlaFB from a cryo EM reconstruction (8.7 Å resolution).** *(A) E. coli* X-ray structure and *A. baumannii* (PDB 6IC4) EM model aligned on MlaB. Inset shows interfaces mapped on both structures. *(B)* Electron density from our crystal structure, corresponding to helices and side chains at the interface between MlaF and MlaB. Sequence alignment of MlaF *(C)* and MlaB *(D)* from the two species

#### Figure S5

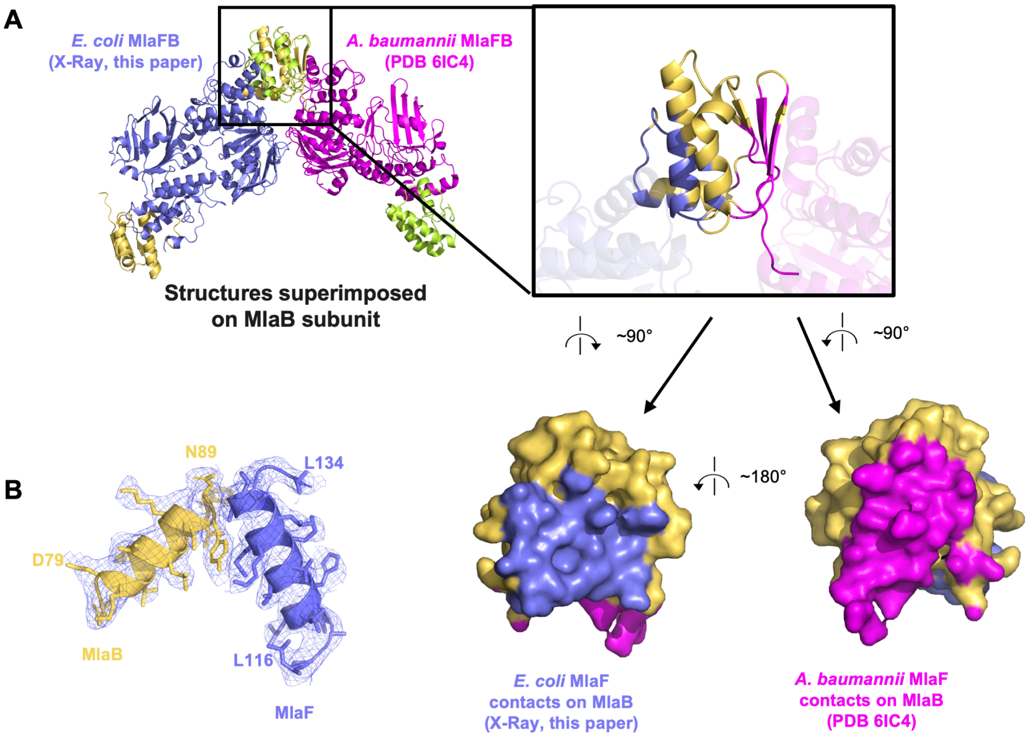


**C**

```
MlaF E. coli          ----MEQSVANLVDMRDVSFTRGNRCIFDNISLTVPRGKITAIMGPSGIGKTTLLRLIGG 56
MlaF A. baumannii     MNNKTPLSTQSLIEVKNLSFNRGERVIYDNISLNIRRGQITAIMGPSGTGKTTLLRLIGG 60
                             *. .*::::::**.**:* *:*****.: **:********* ***********

MlaF E. coli          QIAPDHGEILFDGENIPAMSRSRLYTVRKRMSMLFQSGALFTDMNVFDNVAYPLREHTQL 116
MlaF A. baumannii     QLVPDQGEVLLDGKDIAQMSRQELFAARARMGMLFQSGALFTDMSVYENVAFPIRAHTKL 120
                      *:.**:**:*:**::*  ***..*::.* **.************.*::***:*:* **:*

MlaF E. coli          PAPLLHSTVMMKLEAVGLRGAAKLMPSELSGGMARRAALARAIALEPDLIMFDEPFVGQD 176
MlaF A. baumannii     SENLIAELVALKLESVGLRGTEQLMPTELSGGMNRRVALARAIALDPDLIMYDEPFAGQD 180
                         *: . * :***:*****: :***:****** **.********:*****:****.***

MlaF E. coli          PITMGVLVKLISELNSALGVTCVVVSHDVPEVLSIADHAWILADKKIVAHGSAQALQANP 236
MlaF A. baumannii     PIVKGVLTRLIRSLREALDLTTIIVSHDVPETLSIADYIYVVAEGKIQGEGTPEELQAYA 240
                      **. ***.:** .*..**.:* ::*******.*****: :::*: ** ..*: : ***

MlaF E. coli          DPRVRQFLDGIADGPVPFRYPAGDYHADLLPGS 269
MlaF A. baumannii     SPFVKQFLTGSAEGPVEYQFSHQAYLDNEVRP- 272
                      .* *:*** * *:*** :::    *  : :
```

**D**

```
MlaB E. coli          MSESLSWMQTGDTLALSGELDQDVLLPLWEMREEAVKGIT----CIDLSRVSRVDTGGLA 56
MlaB A. baumannii     ---MVQY--LNQELVVSGKIDFENAEQQYQAGLAIIKKQTSFPLIVDLKQLEHGNTLALA 55
                          :.:   .: *.:**::* :     ::     :*  *     :**.::.: :* .**

MlaB E. coli          LLLHLIDLAKKQGNNVTLQGVNDKVYTLAKLYNLPADVLPR 97
MlaB A. baumannii     VLVQWLRQT-PQKSGLHFKNVPEKMLKIIQACHLQEDLHLV 95
                      :*:: :  :  * ..: ::.* :*: .: :  :*  *:
```

**Supplemental Figure 6. Co-expression of MlaB and MlaF tagged at different positions.**
*(A)* SDS-PAGE of purified His-MlaB alone. *(B)* SDS-PAGE of purified MlaFB complex in which His-tags were placed at different positions as indicated in the figure

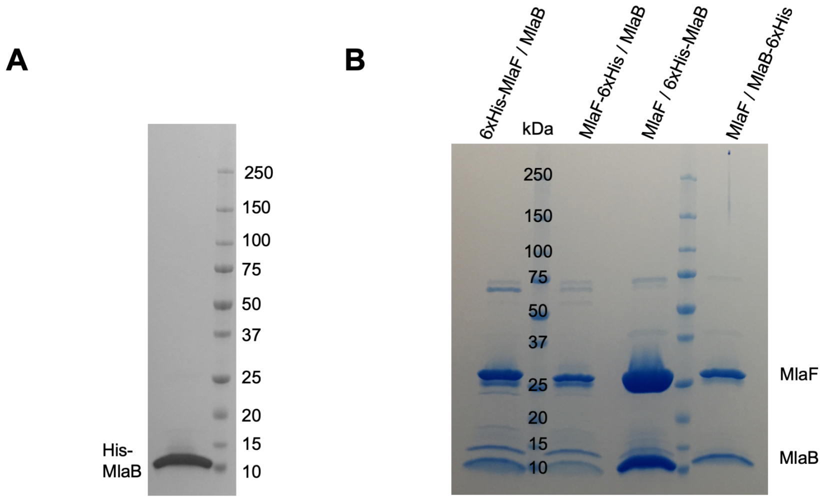

#### Supplementary tables

**Table S1: Bacterial strains.**

| **Strain** | **Relevant Genotype** | **Source** |
|---|---|---|
| **MG1655** | F- lambda- ilvG- rfb-50 rph-1 | Blattner, F. R. et al. |
| **BW25113** | F-, Lambda-, Δ(araD-araB)567, lacZ4787(del)::rrnB-3, rph-1, Δ(rhaD-rhaB)568, hsdR514 | Datsenko, K. A. & Wanner, B. L. |
| **bBEL185** | *mlaB::gfp-FRT* | This work |
| **bBEL190** | *mlaF::FRT* | This work |

#### Table S2: Plasmids

| Plasmid | Relevant Features | Addgene ID | Source |
|---|---|---|---|
| **pBEL1200** | MlaFE-(6xHis-TEV-MlaD)-MlaCB | | Ekiert, *et al*. |
| **pBEL1244** | MlaFE-(6xHis-TEV-MlaD)-MlaC | | This work |
| **pBEL1245** | 6xHis-TEV-MlaF | | This work |
| **pBEL1246** | 6xHis-TEV-MlaB | | This work |
| **pBEL1266** | mlaF, tagless, araBAD promoter | | This work |
| **pBEL1305** | (6xHis-TEV-MlaF)-MlaB | | This work |
| **pBEL1306** | (MlaF-6xHis)-MlaB | | This work |
| **pBEL1307** | MlaF-(6xHis-TEV-MlaB) | | This work |
| **pBEL1308** | MlaF-(MlaB-6xHis) | | This work |
| **pBEL1514** | mlaF(1-246)-mlaE-[6xHis-2xQH-TEV-mlaD]-mlaCB operon | | This work |
| **pBEL1647** | mlaF(Δ247-267), tagless | | This work |
| **pBEL1833** | mlaF(1-250)-GCN4 | | This work |
| **pBEL1840** | MlaF-6xHis | | This work |
| **pBEL1957** | Strep-MlaF-(6xHis-TEV-MlaB) | | This work |
| **pBEL1965** | mlaF(1-246)-(6 aa linker)-GCN4 | | This work |
| **pBEL1966** | mlaF(1-246)-(10 aa linker)-GCN4 | | This work |
| **pBEL1967** | mlaF(1-246)-(14 aa linker)-GCN4 | | This work |
| **pBEL2074** | Strep-MlaF-(6xHis-TEV-MlaB_T52E) | | This work |
| **pBEL2076** | (Strep-MlaF_K47A)-(6xHis-TEV-MlaB) | | This work |
| **pBEL2077** | (Strep-MlaF_E170Q)-(6xHis-TEV-MlaB) | | This work |
| **pBEL2078** | (Strep-MlaF_Y261A)-(6xHis-TEV-MlaB) | | This work |
| **pBEL2079** | (Strep-MlaF_L265A)-(6xHis-TEV-MlaB) | | This work |
| **pBEL2080** | (Strep-MlaF_L266A)-(6xHis-TEV-MlaB) | | This work |
| **pBEL2081** | (Strep-MlaF_Y261A_L265A_L266A)-(6xHis-TEV-MlaB) | | This work |
| **pBEL2082** | (Strep-MlaFΔ[247-269])-(6xHis-TEV-MlaB) | | This work |

**Table S3: Data collection and refinement statistics for crystal structures.**

| | MlaFB (ADP+Mg) | MlaFB (apo) |
|---|---|---|
| **Data collection** | | |
| Space group: | $P3_221$ | $P2_12_12_1$ |
| Cell dimensions: | | |
| a, b, c (Å): | 102.71 102.71 89.71 | 78.27, 136.08, 261.83 |
| α, β, γ (°): | 90, 90, 120 | 90, 90, 90 |
| Resolution (Å): | 44.57-2.9 (3.004-2.9)[1] | 47.8-2.6 (2.693-2.6)[1] |
| Wavelength (Å): | 1.0000 | 1.0000 |
| Observations: | 245,874 | 1,139,469 |
| Unique Reflections: | 12,466 | 86,962 |
| Redundancy: | 19.7 (20.3) | 13.1 (12.6) |
| Completeness (%): | 93.09 (76.44) | 92.07 (75.14) |
| CC1/2: | 0.999 (0.349) | 1.00 (0.538) |
| CC*: | 1.00 (0.72) | 1.00 (0.836) |
| $I/\sigma I$ | 16.09 (1.00) | 18.68 (1.08) |
| $R_{meas}$ | 0.1835 (3.526) | 0.09513 (2.898) |
| **Refinement** | | |
| Resolution (Å): | 44.57 - 2.9 | 48.87-2.60 |
| Reflections (work): | 11,880 | 85,125 |
| Reflections (free): | 586 | 1,834 |
| $R_{work}$ / $R_{free}$ (%): | 19.15 / 24.79 | 19.41/23.21 |
| No. atoms: | | |
| Protein: | 2,760 | 11,361 |
| Water: | 2 | 10 |
| Other: | 29 | 62 |
| Mean B-factor: | | |
| Protein: | 82.75 | 92.48 |
| Water: | 42.31 | 62.94 |
| R.M.S. Deviations: | | |
| Bond lengths (Å): | 0.004 | 0.009 |
| Bond angles (°): | 0.79 | 1.322 |
| Ramachandran plot: | | |
| Favored: | 95.53% | 99.44% |
| Outliers: | 0.84 % | 0.14% |
| Rotamer outliers: | 0.00% | 1.14% |
| Molprobity: | | |
| Molprobity score: | 1.77 | 1.62 |
| Percentile: | 100th | 99th |
| All-atom clashscore: | 8.54 | 11.52 |
| Percentile: | 97th | 96th |
| PDB ID: | TBD[2] | TBD[2] |

[1] Values in parentheses are for highest-resolution shell.

[2] Coordinates are available prior to publication on our website: http://bhabhaekiertlab.org/pdb-links-2
